# Non-contact measurement of emotional and physiological changes in heart rate from a webcam

**DOI:** 10.1101/179812

**Authors:** Christopher R. Madan, Tyler Harrison, Kyle E. Mathewson

**Author notes:** Corresponding author. Email address, Boston College, Department of Psychology, McGuinn 300, 140 Commonwealth Ave., Chestnut Hill, MA, USA 02467.

## Abstract

Heart rate, measured in beats per minute (BPM), can be used as an index of an individual’s physiological state. Each time the heart beats, blood is expelled and travels through the body. This blood flow can be detected in the face using a standard webcam that is able to pick up subtle changes in color that cannot be seen by the naked eye. Due to the light absorption spectrum of blood, we are able to detect differences in the amount of light absorbed by the blood traveling just below the skin (i.e., photoplethysmography). By modulating emotional and physiological stress—i.e., viewing arousing images and sitting vs. standing, respectively—to elicit changes in heart rate, we explored the feasibility of using a webcam as a psychophysiological measurement of autonomic activity. We found a high level of agreement between established physiological measures, electrocardiogram (ECG), and blood pulse oximetry, and heart rate estimates obtained from the webcam. We thus suggest webcams can be used as a non-invasive and readily available method for measuring psychophysiological changes, easily integrated into existing stimulus presentation software and hardware setups.

## Introduction

Heart rate (HR) is a readily measurable index of an individual’s psychophysiological state, specifically autonomic arousal, used in addition to skin conductance response and pupil dilation (Bradley et al., 2008; Kahneman et al., 1969; Robinson et al., 1966). Indeed, the association between the heart and emotional/psychological states dates back to ancient Egypt (Damasio, 1994; Krantz & Falconer, 1997; Schacter & Singer, 1962), as well as permeating into culture throughout the ages (Loe & Edwards, 2004a, 2004b). HR is most often measured using an electrocardiogram (ECG), where changes in voltage generated by innervation of cardiac muscles producing a heartbeat are measured through electrode contacts that are affixed to an individual. However, ECG equipment can be costly, connections can deteriorate over time, and with some participant groups and situations it may be too invasive to apply electrodes. Other less invasive techniques to measure heart rate are therefore needed.

HR can be measured through methods alternative to ECG, such as photoplethysmography (PPG): the detection of variations in transmitted or reflected light (Ackles et al., 1985; Allen, 2007; Jennings et al., 1980; Lu et al., 2009; Schäfer & Vagedes, 2013). Briefly, changes in the light absorbed/reflected by blood can be used to measure the flow of blood. The absorption spectra of blood, and the measurement of the reflectance of skin color in relation to blood, has been studied for many decades within the field of medicine (e.g., Anderson & Parrish, 1981; Angelopoulou, 2001; Brunsting & Sheard, 1929a, 1929b; Edwards & Duntly, 1939; Horecker, 1943; Jakovels et al., 2010, 2011, 2012; Kim & Kim, 2006; Sheard & Brown, 1926; Brunsting & Sheard, 1929; Tsumura et al., 1999, 2000, 2003). A common example of transmission PPG is a pulse oximeter (PulseOx) measurement in hospital settings in which red light is passed through the finger, wrist, or foot and fluctuations in transmitted light are detected.

More recently, a number of studies performed in biomedical engineering laboratories have demonstrated the feasibility of non-contact measuring of HR with a webcam (i.e., a digital video camera that streams its images to a computer). Poh et al. (2010) demonstrated the validity of HR measurements from a webcam by comparing them with measurements obtained at the same time from (but not time synchronized with) a blood pulse oximetry sensor (also see Kwon et al., 2012; Poh et al., 2011a). Subsequent studies have used webcams to study changes in HR due to exercise (Sun et al., 2011, 2012) and the development of devices designed to aid with health monitoring (Poh et al., 2011b; Verkruysse et al., 2008). There have been additional technical advances in how HR is estimated from the webcam recording (e.g., Lewandowska et al., 2011; Pursche et al., 2012; Sun et al., 2012). While these studies have been beneficial in demonstrating the robustness of this approach to measuring HR, the webcam HR estimates were not compared against time-synchronized standard HR measures, and did not evaluate changes in HR as a psychophysiological measure, i.e., the effect of task-related changes on autonomic arousal. As prior studies have indicated lower limits to the sampling rate required to assess ECG signal (Hejjel & Roth, 2004; Pizzuti et al., 1985), it is not clear if the low sampling rate of the webcam will be suitable for measuring heart rate within the context of psychophysiology research.

To test if these techniques could be applied to experimental psychology situations as a method of psychophysiological monitoring, we used a standard webcam to record the light reflected from a participant’s face. Acquisition of HR data from the webcam was marked with respect to events in the stimulus presentation program, which are also marked in concurrently recorded ECG and PulseOx data. While averaging across the face area during recording of the webcam data, to provide anonymity, we measured task-related changes in a participant’s HR. Specifically, we modulated emotional and physiological stress (i.e., viewing arousing images and siting vs. standing, respectively) to elicit changes in HR to demonstrate the use of a webcam as a psychophysiological measurement of autonomic activity.

As a first test of event-related physiological changes in HR, we measured HR in a blocked sitting vs. standing task where we expected to observe large within-subject, task-related differences in HR. HR was measured concurrently from participants using the webcam along with ECG and pulse oximetry, for comparison. Briefly, when standing, the heart has to work harder to pump blood to the extremities to ensure sufficient force to overcome the effects of gravity (Caro et al., 1978; Herman, 2007; Rushmer, 1976). Empirically, the difference in HR for sitting vs. standing is approximately 8-10 BPM in young adults (Guy, 1837; MacWilliam, 1933; Schneider & Truesdell, 1922; also see Stein et al., 1966).

As a test of the feasibility of webcam HR in a task-related context, we next measured changes in HR time locked to emotional pictures, again concurrently with all three measures. Within the literature on emotional processing (e.g., Bradley et al., 2001a, 2008; Buchanan et al., 2006; Critchley et al., 2013; Garfinkel & Critchley, 2016; Lang et al., 1993; Levenson, 2003), it is well known that viewing emotionally arousing stimuli increases autonomic arousal, across a variety of psychophysiological measures. Presentation of unpleasant (i.e., negative valence) pictures elicits a deceleration in HR, referred to as fear bradycardia, and that this deceleration is primarily mediated by the autonomic/parasympathetic nervous system (Bradley et al., 2001a, 2001b; Campbell et al., 1997). Hare (1973) suggested that this HR deceleration could be due to an orienting response, rather than a defensive response, to viewing the picture (also see Graham & Clifton, 1966; Sokolov, 1963). Empirically, this deceleration is a change of approximately 1-3 beats per minute (BPM), with a time course of approximately 6 seconds (Abercrombie et al., 2008; Bradley et al., 2008; Buchanan et al., 2006; Hare, 1973). Here we tested if our webcam HR technique would provide sufficient sensitivity to measure the subtle changes associated with a typical psychophysiological experiment, with the ECG and pulse oximetry data also acquired for comparison.

## Method

### Participants

A total of 24 volunteers participated in the experiment (age: *M*=21.7, range=18-25; 14 female) and were recruited from the University of Alberta community using advertisements around campus. Sample size was determined based on pilot studies of the sitting vs. standing task. All participants gave informed, written consent and were compensated at a rate of $10/hr for their time. The experimental procedures were approved by an internal research ethics board of the University of Alberta.

### Equipment

Video was recorded using a Logitech HD Pro Webcam C920 (Logitech International S.A., Newark, CA). The webcam video was recorded in color at a resolution of 640×480, at a mean sampling rate of 12 Hz (0.083±0.016 s [*M±SD*] between video frames). Stimuli were presented on a Dell UltraSharp 24” monitor with a resolution of 1920×1200, using a Windows 7 PC running MATLAB R2012b (The MathWorks Inc., Natick, MA) with the Psychophysics Toolbox v. 3 (Brainard, 1997). Webcam data was simultaneously recorded using in-house code in the same MATLAB script as the stimulus presentation.

ECG signals were collected from bilateral wrists of participants using Ag/AgCl snap-type disposable hydrogel monitoring electrodes (ElectroTrace ET101, Jason Inc., Huntington Beach, CA) in a bi-polar arrangement over the distal extent of the flexor digitorum superficialis muscle, with a ground over the distal extent of the left flexor carpi radialis. Prior to applying the electrodes, the participant’s skin was cleaned using alcohol wipes. Blood pulse oximetry data was collected using a finger pulse sensor attached to the index finger of the participant’s right hand and enclosed in a black light blocking sheath (Becker Meditec, Karlsruhe, Germany). Both sensors were connected to the AUX ports of a BrainVision V-Amp 16-channel amplifier (Brain Products GmbH, Gilching, Germany) using BIP2AUX converters. Physiological data was recorded at 500 Hz at 1.19 μV/bit using BrainVision Recorder software (Brain Products GmbH) with a band-pass online filter between 0.628 and 30 Hz.

For the ECG and pulse oximetry data, data was collected for the entire duration of each task (sit-stand, emotion). In order to mark the time of stimulus onset in the ECG and pulse oximetry data, an 8-bit TTL pulse was sent via parallel port by the stimulus presentation software coincident with the onset of important stimuli, marking their time and identity (i.e., onset/offset of the fixation and pictures). The webcam data was recorded in epochs for each block (in the sit-stand task) or trial (in the emotion task) by the stimulus presentation software yoked to the stimulus display. The task presentation and the data collection through all three measures were done by the same computer, allowing for all signals to be easily synchronized.

### Stimuli

The pictures selected for the emotion task comprised four categories, each with 15 pictures/category. The pictures were selected from the International Affective Picture System (IAPS; Lang et al., 2008) database based on normative ratings for valence and arousal and were supplemented with pictures used in prior studies of emotional processing (Singhal et al., 2012; Wang et al., 2005, 2008). Mean IAPS valence/arousal scores (9-point scale, as described below) of the four categories were as follows: Neutral (Neut; 5.8/1.6), Low Arousal (Low; 3.6/3.3), Medium Arousal (Med; 2.3/5.8), and High Arousal (High; 2.3/6.1). A repeated-measures ANOVA showed that valence ratings for each category were significantly different from each adjacent category except for Med and High (i.e., Neut > Low > Med = High, [*F*(3,72) = 132.97, *p* < .001]). A repeated-measures ANOVA of arousal ratings showed that each category was significantly different from each adjacent category such that, Neut < Low < Med < High [*F*(3,72) = 150.59, *p* < .001]. Pair-wise comparisons were Holm-Bonferroni-corrected.

### Procedure

The experiment was conducted in a room of an experimental lab with normal lighting conditions. The experiment consisted of two tasks: blocks of sitting and standing (sit-stand task), and passive viewing of emotional and neutral pictures (emotion task). Task order was pseudorandomized across participants. In both cases, participants were seated in front of a webcam, which was placed either on a tripod (sit-stand task) or on top of the computer monitor (emotion task).

***Sitting vs. standing task***. The sit-stand task contained 10 blocks, of 30 s each. In half of the blocks, participants were instructed to be seated, in the other half they were to stand. The order of the blocks was pseudorandomized such that no more than two blocks from the same condition (e.g., sitting) occurred sequentially.

Before each block, the tripod was adjusted to suit the participant’s height. The participant was then instructed to be as still as possible during the 30 s of data collection. ***Emotional and neutral picture-viewing task.*** The emotion task was comprised of three blocks, each consisting of 20 trials. On each trial, participants were first shown a scrambled picture with a fixation cross (“+”) overlaid, followed by an emotional or neutral picture, then followed by the scrambled picture again. Pictures were presented for 2000 ms; scrambled stimuli were presented before and after each picture for 500 and 3000 ms, respectively. The scrambled stimuli were scrambled versions of the emotional or neutral picture, converted to grayscale and kept isoluminant with the picture. The order that the pictures were presented was pseudorandomized such that no more than two stimuli from the same category (e.g., high arousal) were shown sequentially. Trials were separated by jittered inter-trial intervals, ranging from 5000 to 6500 ms.

Prior to each block, the webcam recording was calibrated such that the participant aligned their head with a template indicating the area-of-interest (AOI) using live video feedback. Once the AOI was sufficiently aligned with the participant’s face, they were instructed to place their hands on the table in front of them and to remain as still as possible while the stimuli were presented and data was recorded.

### Data Analysis

The processing workflow for the webcam analyses is outlined in Figure 1. Based on the calibration, a rectangular AOI positioned over the participant’s face constrains the collection of the webcam data. To ensure the collected data preserved participant anonymity, color values for each frame were averaged across this AOI during data collection, rather than maintaining the raw webcam frame. As a result, we only retained three intensity values per webcam frame, corresponding to red, green, and blue (RGB) channels. Data for each block (sit-stand task) or trial (emotion task) were then saved for offline analyses.

**Figure 1.**
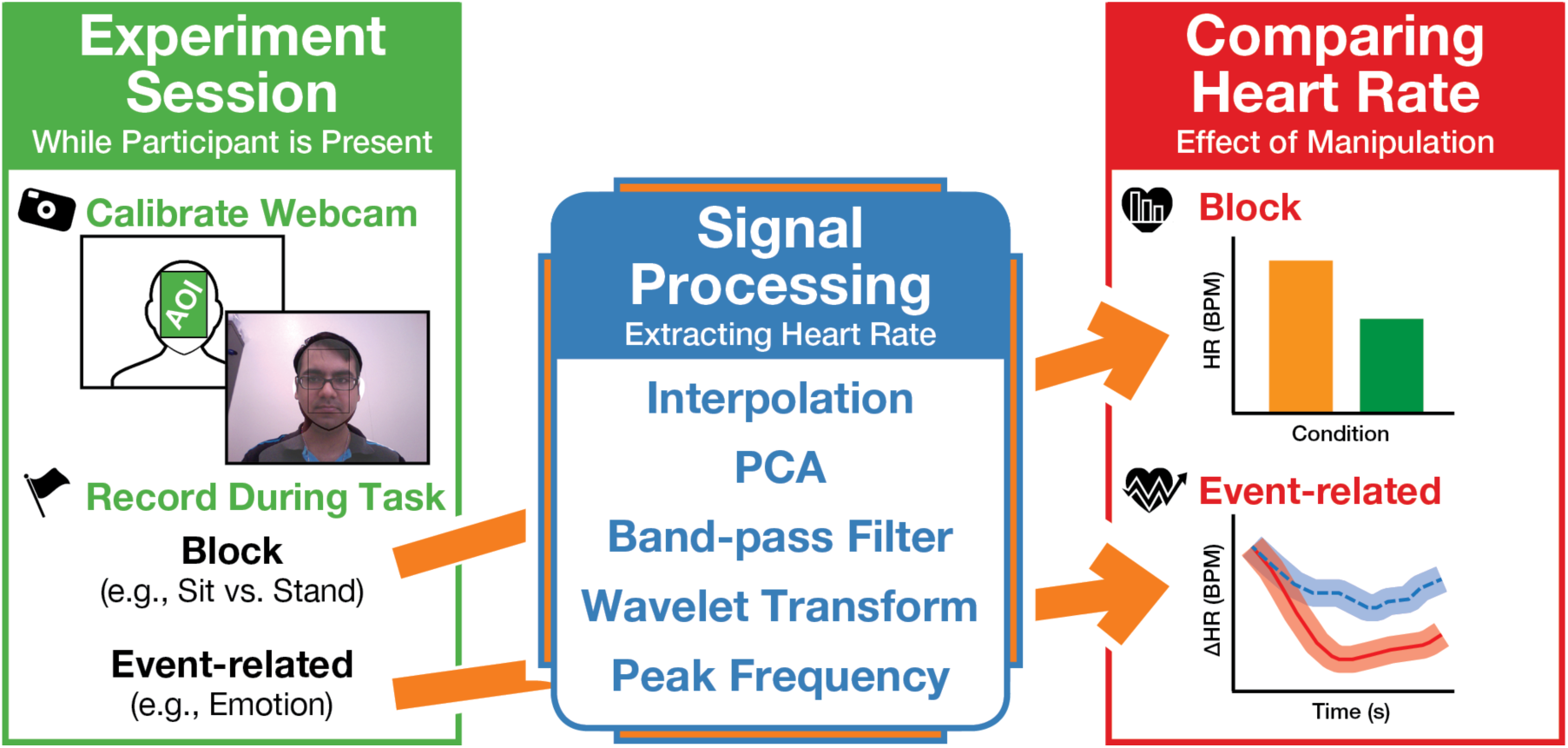
Illustration of the analysis pipeline.

Three pre-processing steps were used specifically on the continuous webcam data from entire blocks. First, to maximize the temporal resolution of the webcam data, we had sampled frames from the webcam as quickly as the hardware would allow (using the `videoinput` function in MATLAB), which lead to a non-uniform sampling rate. As minor fluctuations in the interval between successive frames would influence our estimated heart rate, we re-sampled the webcam data with a uniform interpolation of 12 Hz using the `interp` function in MATLAB. As a note to other researchers, if your hardware is able to sample from the webcam at a higher rate reliably, it would be simpler to instead have a uniform sampling rate and not necessitate re-sampling via interpolation. Second, it has been demonstrated that the green RGB color channel is the most sensitive to changes in light reflectance associated with oxygenated vs. deoxygenated blood, though the red and blue channels do still contain plethysmographic information (Lee et al., 2013; Poh et al., 2010; Sun et al., 2011, 2012; Verkruysse et al., 2008). To maximize info from all channels, we submitted the three color-channel time-series data (for the entire block) into a principal component analysis (PCA), allowing us to extract the variability in signal that was common across the three channels. We used the coefficients from the second principal component as our time-series data, as this was the component that corresponded to HR-related changes in all cases (also see Lewandowska et al., 2011; Poh et al., 2010, 2011a, 2011b; Pursche et al., 2012; Tsumura et al., 2000). Third, an additional offline Butterworth band-pass filter was applied to the data (high=0.8 Hz, low=3.0 Hz; see Gribok et al., 2011). This provided a 12-Hz signal from the webcam continuous throughout each block, along with the 500 Hz signals from the ECG and PulseOx.

Finally, for each each measure (webcam, ECG, PulseOx), the continuous data at submitted to a continuous wavelet (Morlet) transform implemented in the BOSC library (“Better OSCillation detection”; Hughes et al., 2012; Whitten et al., 2011). The transform was used to obtain the power spectra for the frequencies corresponding to a range of plausible heart rates, 50-140 BPM, in 1 BPM increments, and a wavelet number of 6. At each time point of the resulting spectrogram, heart rate was calculated as the frequency with the highest power.

#### Blocked design

For the sitting vs. standing task, heart rate was estimated as a single value for each trial. Heart rate for each trial, for each measure, was estimated as the median heart rate for the 30-s block.

#### Event-related design

For the emotional and neutral picture-viewing task, heart rate was measured as a time-varying change, in relation to the onset of the image. To compute the event-related variations in HR, changes in HR were estimated using a sliding time-window. For each trial, epochs spanning from 5 s before to 5 s after the onset of the picture, were segmented from the continuous data.

Preliminary analyses indicated that the webcam data was confounded by stimulus luminance, where the luminance of the presented picture would interact with photoplethysmography signal intended to be recorded. This occurred despite pictures being preceded by an isoluminant scrambled picture; this likely occurred because trial-wise differences in the light emitted by the monitor when presenting the pictures influenced the light reflected by the participants’ face and detected by the webcam. To address this confound, luminance for the pictures was regressed out of the individual trial timecourses. Luminance here was quantified by converting the pictures to CIELab 1976 color space, and summarized as a single value for each picture by averaging across the L* channel. For future research, we recommend matching the stimulus luminance across pictures if possible, making this regression step unnecessary. The presentation of the scrambled picture is critical, however, to prevent changes in screen luminance that correspond to the onset and offset of the picture-of-interest. We also recommend the scrambled picture be presented in grayscale as color properties of the original pictures may not be matched across conditions (e.g., high arousing pictures were more red than neutral pictures).

For each trial and measure, the average heart rate in the 2000 ms prior to the picture onset was then subtracted from the entire trial period to align the picture onset across trials, i.e. a baseline correction. Then, for each HR recording type, separate averages are created for each subject in each of the emotional picture conditions. For statistical tests, the peak deceleration between 1500 and 3000 ms was used (based on prior findings; e.g., Abercrombie et al., 2008; Bradley et al., 2008; Buchanan et al., 2006), measured for each participant and emotion condition. See Figure 2 for a demonstration of the analysis pipeline for an event-related design.

**Figure 2.**
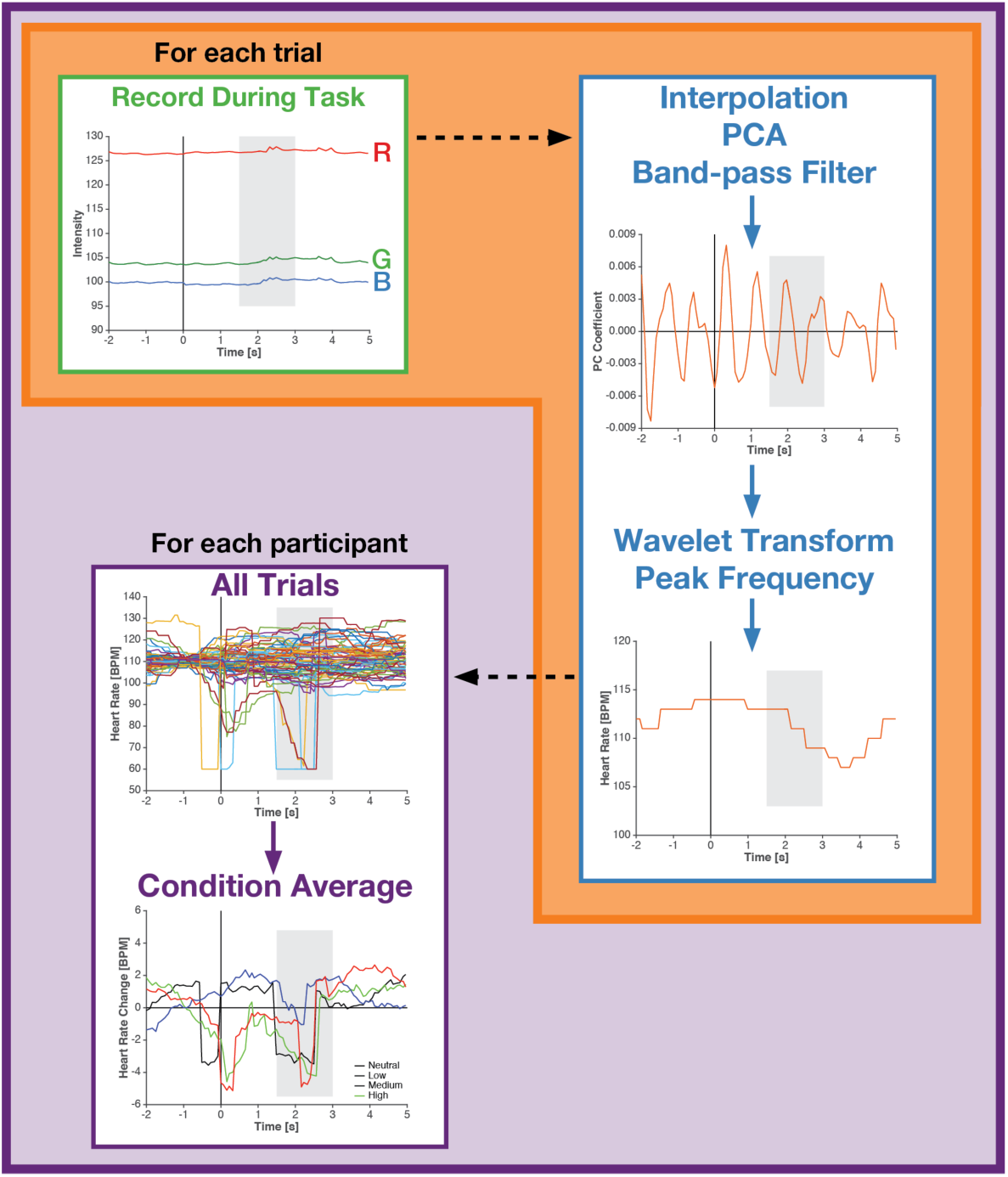
Demonstration of the analysis pipeline for an event-related design.

#### Data quality

To ensure that the heart rate estimates obtained from the ECG and PulseOx data were sufficiently reliable, we excluded participants where the power at the peak frequency was less than twice the mean power in the sitting vs. standing task (*N*=1). ANOVA results are reported with Greenhouse-Geisser correction for non-sphericity where appropriate.

## Results

### Sitting vs. standing task

We first compared heart rate measurements for sitting vs. standing with each measurement method using a 2 [*Posture*: Sit, Stand] × 3 [*Measure*: ECG, Pulse Oximetry (PulseOx), Webcam] repeated-measures ANOVA, averaging across block. As shown in Figure 3A, we observed a main effect of Posture [*F*(1,22)=85.29, *p*<0.001, *η*_*p*_^2^=0.80], where standing was associated with a 10.4 BPM increase in heart rate relative to sitting. Neither the main effect of Measure [*F*(1,28)=2.29, *p*=0.14, *η*_*p*_^2^=0.09] nor the interaction [*F*(2,42)=0.15, *p*=0.85, *η*_*p*_^2^=0.007] were significant. Planned contrasts showed that the effect of posture was observable using each measure individually [ECG: *t*(22)=8.92. *p*<0.001, Cohen’s *d*=0.82, *M*_*diff*_ = 10.5 BPM; PulseOx: *t*(22)=7.84, *p*<0.001, *d*=0.82, *M*_*diff*_ = 10.6 BPM; Webcam: *t*(22)=9.41, *p*<0.001, *d*=0.90, *M*_*diff*_ = 10.2 BPM].

**Figure 3.**
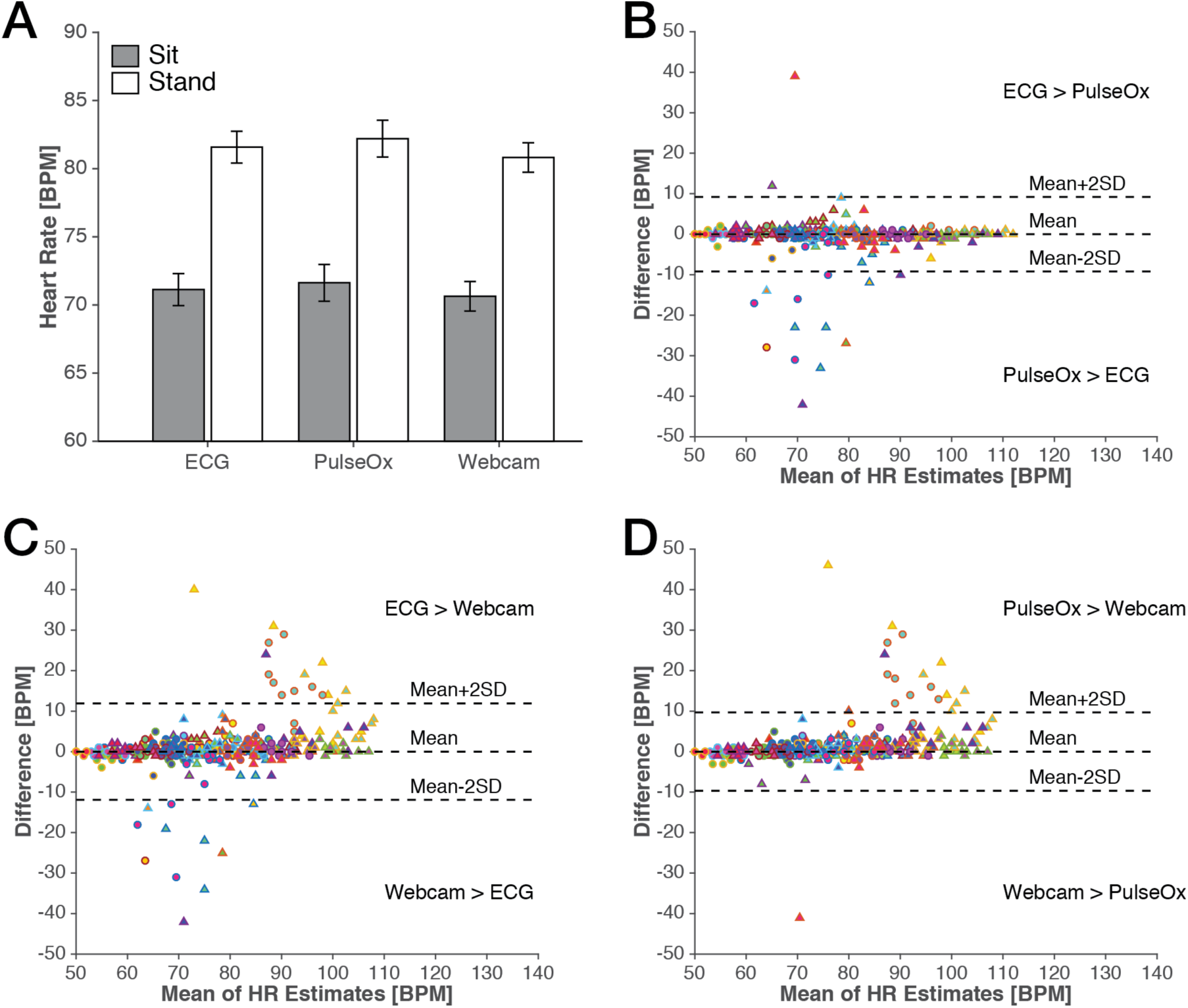
Results from the sitting vs. standing task. (A) Mean heart rate for sitting and standing from each measure. Error bars represent SEM, corrected for inter-individual differences (within-subject SEM; Loftus & Masson, 1999). Bland-Altmann plots for pairs of measures: (B) ECG-PulseOx, (C) ECG-Webcam, and (D) PulseOx-Webcam. Markers represent each block of the task from each participant. Markers in distinct colors represent individual participants; measurements from sitting blocks are shown as circles, standing blocks are shown as triangles.

To evaluate the agreement between the measurements more precisely, we additionally compared the heart-rate estimates from each block, i.e., 10 measurements per participant, between the three measures using correlations and Bland-Altman analyses. All three pairwise correlations were high and of similar magnitude [ECG–PulseOx: *r*(458)=0.950; ECG–Webcam: *r*(458)=0.913; PulseOx–Webcam: *r*(458)=0.944], as were the concordance correlation coefficients (Lin, 1989) [ECG–PulseOx: *r*(458)=0.949; ECG–Webcam: *r*(458)=0.907; PulseOx–Webcam: *r*(458)=0.935]. In all three cases, 2 SD of the difference between the compared measurements was approximately 10 BPM, as shown in Figures 3B-D [ECG–PulseOx: 9.19 BPM; ECG–Webcam: 11.91 BPM; PulseOx–Webcam: 9.67 BPM]. We did, however, observe a greater degree of bias when using the webcam, relative to the other measurements [ECG-PulseOx: –0.56 BPM; ECG– Webcam: 0.63 BPM; PulseOx–Webcam: 1.19 BPM]. This bias suggests that the webcam tends to slightly underestimate heart-rate estimates, perhaps due to the increased noise or slower sampling rate of the webcam measurement. Moreover, considering that certain participants are overrepresented in the outliers it is likely the case that some artifactual noise was impairing the ability to reliability determine the heart rate using some of the measures for these individuals. For instance, hair or clothes, as well as makeup, could interfere with the webcam measurement leading to unrealiable estimates of HR on those blocks.

### Emotional and neutral picture-viewing task

As shown in Figure 4A-C the heart-rate decelerations for several of the conditions did not differ. Using the same stimuli in an fMRI study, Hrybouski et al. (2016) found that medium and high arousal stimuli were not distinct in behavioural ratings of emotional arousal or amygdala fMRI (BOLD) activity, and thus collapsed them together in their reported analyses. Similarly, to maximally index the effect of the emotional pictures on heart rate, here we examined the mean response to the high and medium arousal picture conditions, compared to both the pre-stimulus baseline or viewing of the neutral pictures (Figure 3D). Thus, we pooled high and medium arousal images together and dropping the low arousal condition, as done in Hrybouski et al. (2016), as shown in Figure 4D-F.

**Figure 4.**
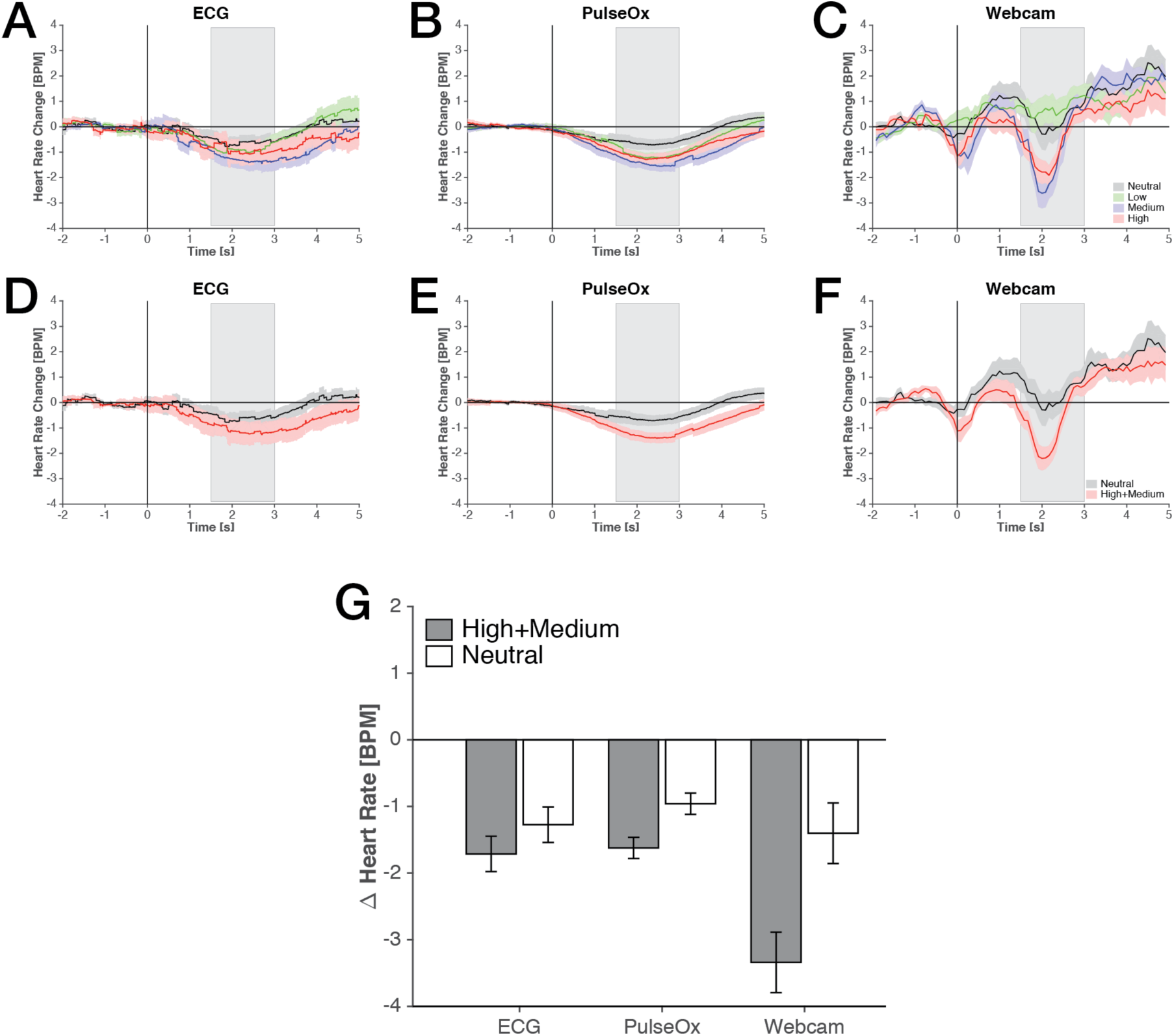
Results from the emotional and neutral picture-viewing task. Event-related changes in heart rate in response to viewing each of the picture types, as measured by the (A) ECG, (B) PulseOx, and (C) Webcam. Shaded error bars represent within-subject SEM. The shaded time window (1500-3000 ms) depicts the data used in the statistical analyses. (D-F) Re-plots panels A-C, collapsing the High and Medium arousal conditions and removing the Low arousal condition. (G) Mean heart rate deceleration related to stimulus presentation, relative to the pre-stimulus baseline. Error bars represent SEM, corrected for inter-individual differences (within-subject SEM; Loftus & Masson, 1999).

We examine the heart-rate deceleration effects using a 2 [*Emotion*: High+Medium, Neutral] × 3 [*Measure*: ECG, Pulse Oximetry (PulseOx), Webcam] repeated-measures ANOVA, based on the mean heart rate during the analyzed window between 1500 and 3000 ms, relative to the pre-stimulus baseline (see Figure 4G). We observed a main effect of Emotion [*F*(1,22)=7.94, *p*=0.010, *η*_*p*_^2^=0.23], where the High+Medium pictures were associated with a 1.01 BPM decrease in heart rate relative to Neutral pictures. Neither the main effect of Measure [*F*(1,23)=2.58, *p*=0.12, *η*_*p*_^2^=0.11] nor the interaction [*F*(1,24)=1.56, *p*=0.22, *η*_*p*_^2^=0.068] were significant.

Despite the non-significant interaction, as planned contrasts we nonetheless report the HR effects for each measure. With the ECG data we observed a significant heart-rate deceleration of 1.71 BPM relative to the pre-stimulus baseline [*t*(22)=4.40, *p*<0.001, *d*=0.96], as well as a nominal deceleration of 0.44 BPM relative to viewing neutral pictures in the same window [*t*(22)=0.83, *p*=0.42, *d*=0.28]. The pulse oximetry data presented similar effects of viewing the emotional stimuli [relative to baseline: *t*(22)=4.81, *p*<0.001, *d*=1.04, 1.62 BPM deceleration; relative to neutral pictures: *t*(22)=2.08, *p*=0.049, *d*=0.52, 0.66 BPM deceleration]. With the webcam we observed a significant heart-rate deceleration of 3.33 BPM relative to the pre-stimulus baseline [*t*(22)=4.37, *p*<0.001, *d*=0.95], as well as a deceleration of 1.94 BPM relative to viewing neutral pictures in the same window [*t*(22)=2.14, *p*=0.044, *d*=0.57]. Thus, we observed significant heart-rate decelerations for emotional pictures with the pulse oximetry and webcam measures, but not with ECG. While the ECG and pulse oximetry obtained similar decelerations due to the arousing pictures, the ECG measure had slightly more variance in the effect (see Figures 4D and E).

It is not clear why the webcam is yielding pronounced, and narrower, heart-rate deceleration effects, particularly since it has less temporal resolution than the other two measures. It is possible that the webcam is measuring autonomic changes in addition to those related to photoplethysmography, such as effects of temperature (influencing skin vasculature) or face-specific responses such as emotion-related changes in facial expressions or blushing. Vasoconstrictive or vasodilative changes associated with sympathetic activity may have also contributed. Future research is needed to better understand how these other factors can influence HR estimates obtained from face recordings. These additional factors may also be responsible for the slight acceleration detected just prior to the deceleration (i.e., the peak at approximately 0.75s in Figure 4F).

## Discussion

Heart rate can change in relation to psychological processes, in addition to physiological states. Here we demonstrated that a standard webcam can readily be used as a heart rate measurement device. Despite limitations in sampling rate, we were able to measure small heart-rate decelerations commonly associated with processing emotional pictures, in addition to the much larger changes in heart rate that are known to be associated with physiological state changes.

Our results showed very close agreement with conventional techniques measured simultaneously in both blocked and event-related designs. Differences in the webcam in the block design could largely be attributed to two outlier subjects for whom the webcam reliably underestimated their heart rate (HR). Therefore some individuals seem to be better conceal from the camera their on-going HR. We cannot investigate in the current data set further to determine what characteristics physically or behaviourally were associated with these imprecisions (e.g., we only saved the webcam data for the face AOI, not the full webcam frame; did not collect inter-individual difference measures), but future work should better understand such individual differences in the measurement success.

Measuring non-contact physiological changes in HR over long periods of time as we showed in our sit-stand results provides an important tool by which one could, in real time, or on recorded footage, identify the ongoing HR of individuals under various levels of physical activity, or in various situations. The live video itself can even be modified to accentuate or visualize the pulse and heart rate on the body (Poh et al., 2011a).

The work here was intended to serve as a proof-of-principle that measurement of HR via webcam is sensitive enough for psychological studies. HR decelerations have been shown to index subsequent memory (Abercrombie et al., 2008; Buchanan et al., 2006; Cunningham et al., 2014; Fiacconi et al., 2016; Garfinkel et al., 2013; Jennings & Hall, 1980), task difficulty (Kahneman et al., 1969), introceptive awareness (Garfinkel et al., 2013), and state anxiety (Garfinkel et al., 2014; Schachter & Singer, 1962). Heart rate is also known to be coupled to other physiological measures such as pupil dilation, skin conductance, and microsaccades (Bradley et al., 2008; Kahneman et al., 1969; Ohl et al., 2016). Consideration is needed to determine the applicability of this webcam approach, however, as it may not be suitable sensor of heart rate in all cases. For instance, heart-rate variability (HRV) has been associated with physiological well-being, and is related to a variety of factors including autonomic regulation and reactivity to acute stressors (e.g., Francis et al., 2015; Hallman et al., 2011; Shaffer et al., 2014). However, the current sampling rate of 12 Hz is insufficient, where HRV usually requires a sampling rate of 250 Hz or higher (Hejjel & Roth, 2004; Pizzuti et al., 1985; Schäfer & Vagedes, 2013). Higher-end webcams or other video cameras, i.e., high-speed cameras, may be able to acquire data at a suitable sampling rate for HRV analyses, though testing will be necessary to determine other limiting factors, such as the rate of MATLAB’s video I/O protocol. Further research is also necessary to establish the boundary conditions or other hardware limitations associated with future applications of this webcam approach to measuring HR, such as an index of vasculature function.

From a technical standpoint, measuring heart rate using a webcam can afford several benefits relative to the standard approaches such as ECG and pulse oximetry. While these other measures are non-invasive, a webcam is additionally non-contact. Thus, a webcam can be used equally well with participants that may have sensitive or delicate skin, such as older adults or patient populations, where contact measurements may be problematic. Furthermore, the impedance of the connection between the ECG electrode and the skin may increase over time leading to increased noise in ECG HR estimates. Pulse oximetry can similarly become dislodged over time due to its placement on the finger, and is cumbersome and interferes with normal typing and movements. Webcam equipment is also much more available and affordable than ECG and pulse oximetry, potentially making heart rate analyses more cost effective for pilot studies or researchers with limited funding.

A webcam may also used to covertly measure heart rate with the participant being unaware that this data is even being collected, as long as proper consent and IRB protocols are followed. For instance, covert heart-rate recording could be beneficial along with a Concealed Information Test (see Matsuda et al., 2012, for a review). In this case, it is additionally useful to point out that the webcam need not be calibrated towards the participants’ face, but merely needs to record video data from exposed skin, e.g., an arm, in the presence of sufficient ambient lighting. Others have previously demonstrated that a single webcam can be used to measure heart rate for several individuals simultaneously (Poh et al., 2010). Additionally, the use of webcams to measure heart rate could be beneficial to medical care, such as when using video communication in patient care (see Armfield et al., 2012). Although animals may seem like unlikely candidates for such measurement, the exposed skin on the face and ears of mammals can also provide a non-invasive window into single or multiple animal HR monitoring.

One could argue that the usefulness of this technique is limited by the requirement of the subject to be still in the camera focus. Others have circumvented by using face detection algorithms (Poh et al., 2010, 2011b) or could take advantage of signal filters designed for detecting skin pigments (Anderson & Parrish, 1981; Changizi et al., 2006; Edwards & Duntly, 1939; Tsumura et al., 1999, 2003). If desired, multiple cameras and 3D motion trackers could be used to improve face/skin localization. Furthermore, movement artifacts are a similar problem for both ECG and PulseOx measurement. For experiment implementation, here we used the Psychophysics Toolbox and MATLAB. Functions within the Psychophysics Toolbox were used to present the stimuli while base MATLAB functions were used to interface with the webcam hardware. This allowed us to yolk webcam data recording to the stimulus presentation, but future studies could further integrate presentation and webcam recording for use with biofeedback (also see Lakens, 2013). In sum, here we demonstrated that the webcam is sufficiently sensitive for psychologically relevant changes in heart rate, opening many potential lines of future research.

## Acknowledgements

This work was supported by a NSERC discovery grant and startup funds from the Faculty of Science to KEM. CRM was supported by a fellowship from the Canadian Institutes of Health Research (FRN-146793).

